# HnRNPA2B1 tunes antimycobacterial immune responses in macrophages through alternative splicing of *Irgm1*

**DOI:** 10.64898/2025.12.16.694586

**Authors:** MJ Chapman, JB Huskey, KS Armijo, S Hahn, J Spellman-Reliford, AK Coleman, CJ Mabry, S Carpenter, RO Watson, KL Patrick

## Abstract

Onset and progression of active tuberculosis disease result from upsetting the delicate balance between Mtb virulence and host defenses. Because it dynamically tunes the functional output of protein expression in cells, alternative splicing, a process by which different mRNAs can be generated from a single gene, is positioned to play a critical role in maintaining an equilibrated Mtb-macrophage host-pathogen interface. To gain insight into how alternative splicing shapes anti-mycobacterial immune responses, we used RNA-sequencing and splicing-aware computational pipelines to quantify alternative splicing in Mtb-infected bone marrow-derived murine macrophages. We found that ∼5% of expressed macrophage genes exhibit one or more splicing changes at 8h post-Mtb infection, highlighting alternative splicing as a key regulatory node in the macrophage response to Mtb. We next sought to identify RNA binding proteins that play an out-sized role in shaping the macrophage transcriptome during Mtb infection. We discovered that the splicing factor heterogeneous nuclear ribonucleoprotein A2B1 (hnRNPA2B1) promotes the early induction of inflammatory genes while dampening several type I interferon-stimulated genes in response to Mtb. HnRNPA2B1 also controls alternative splicing of many genes during Mtb infection, including Irgm1, a critical immunity-related GTPase. The balance of Irgm1-long vs. -short is differentially regulated in response to diverse inflammatory cues and macrophages overexpressing Irgm1-short are defective in autophagosomal targeting, lysosomal homeostasis, and restriction of Mtb replication. These data highlight a key role for AS in shaping the macrophage transcriptome and pinpoint hnRNPA2B1 as a novel restriction factor in the cell-intrinsic response to Mtb.

**IMPORTANCE:** Although the process of making proteins from RNAs requires many steps (transcription, capping/polyadenylation, pre-mRNA splicing, mRNA export, mRNA modifications, etc.), we know very little about how post-transcriptional steps contribute to host immune defenses. Here, we show that alternative splicing, the process of making different mature RNAs from a single precursor RNA, is a prominent and dynamic feature of macrophage infection with the bacterial pathogen *Mycobacterium tuberculosis* (Mtb). We identify the splicing regulator hnRNPA2B1 as a key coordinator of early gene expression during Mtb infection, influencing pathways that promote inflammation and help restrict bacterial growth. Notably, we report that hnRNPA2B1 controls the splicing of the antimycobacterial protein *Irgm1* to generate different flavors of the protein. Since only one Irgm1 flavor can restrict Mtb growth inside macrophages, maintaining the balance of these proteins in response to diverse inflammatory cues is important. By revealing how RNA processing shapes the macrophage response to Mtb, our work highlights an often-overlooked layer of immune regulation and opens new avenues for splicing-targeted therapies designed to boost Mtb killing in macrophages.

## INTRODUCTION

Upon sensing pathogens like *Mycobacterium tuberculosis* (Mtb) macrophages rapidly reprogram their transcriptomes to engage antimicrobial defenses and alert neighboring cells to a newly detected threat. This response is initiated by Pattern Recognition Receptors (PRRs), which detect specific Pattern Associated Molecular Patterns (PAMPs) in distinct subcellular compartments. PAMP recognition by PRRs triggers signaling cascades that activate transcription factors such as IRF3, AP-1, and NF-kB, which in turn, drive rapid induction of immune response genes (1). The flexibility and specialization of PRR signaling allows macrophages to tailor responses to different pathogens. Mtb is sensed by multiple PRRs at different stages of infection, with TLR1/2/6 sensing of Mtb outer membrane glycoproteins and mycolic acids occurring at the plasma membrane and cGAS sensing of Mtb dsDNA occurring in the cytosol, following destabilization of the Mtb-containing phagosome (2–4). Together, these compartment-specific sensing pathways allow macrophages to integrate multiple microbial cues and calibrate their antimicrobial programs.

In addition to activation of transcription, post-transcriptional regulatory steps such as 3’end formation and polyadenylation (5, 6), mRNA decay (7–10), and pre-mRNA splicing (11–18) are increasingly appreciated as critical determinants in shaping the innate immune milieu. Among these, splicing stands out because of its ubiquity and potential for diversifying the proteome. In the murine and human genomes, ∼90% of protein-coding genes contain multiple exons (19) and ∼95% of multi-exon genes are predicted to undergo alternative splicing (AS) (20, 21), a process whereby exons are selectively included or excluded in a regulated manner (22, 23). Through AS, a single gene can generate distinct isoforms from a common pre-mRNA, thereby expanding proteomic diversity independently of transcriptional activity (24–27).

Splicing decisions (where and when to splice) are controlled by RNA binding proteins, particularly factors in the SR (serine-arginine rich) and hnRNP (heterogenous nuclear ribonucleoprotein) families (22, 23). Generally, SR proteins promote splicing and exon inclusion while hnRNPs inhibit splicing, causing exon skipping. SRs and hnRNPs have gained recent attention as orchestrators of immune cell reprogramming (11, 12, 16, 28–31). In macrophages, hnRNP M negatively regulates abundance of inflammatory transcripts like *Il6* by slowing intron removal (12) and SRSF6 limits tonic interferon signaling by controlling alternative splicing of the mitochondrial pore-forming protein Bax (11). Splicing factors also have emerging roles in T cell immunity, where the SR protein Tra2b regulates *TCR* transcript splicing to decide T cell fate (28) and SRSF1 regulates splicing of *Cd6*, a protein involved in coordination of antigen presentation (32). Together, these studies support a model whereby immune cells rely on splicing factors to execute specific responses to pathogens and/or inflammatory cues.

Earlier studies have established that Mtb infection elicits hundreds of AS events in human macrophages in functionally diverse genes including *Il12rb1* (involved in dendritic cell migration), *Rab8b* (important for phagolysosomal maturation), and *Pgk1* and *Acsl1* (genes involved glycolysis and lipid synthesis) (33, 34). AS events have also been identified as potential biomarkers for tuberculosis infection in human sera (35). Mechanistically, the inputs that influence AS during Mtb infection are likely multifactorial, with growing evidence that immune stimuli and Mtb effector proteins can influence splice site selection (11, 36–39).

We recently became interested in a splicing factor called hnRNPA2B1. HnRNPA2B1 is a hnRNP A/B family member closely related to hnRNPA1 (40). HnRNPA2B1 has 4 annotated domains; an RNA binding domain composed of two tandem RNA Recognition Motifs, an RGG box, a low complexity prion-like domain, and a PY-motif containing a M9 nuclear localization signal (PY-NLS) (41). HnRNPA2B1 plays known roles in neurologic development (42) and cancer (43–46), but remains understudied in the context of immunity. Outside of its role in splicing, hnRNPA2B1 can also act as a reader of mRNAs methylated by METTL3 (47–49), shuttle microRNAs in extra-cellular vesicles (50, 51), and protect transcripts from degradation (52, 53). Leveraging an hnRNPA2B1 conditional knockout mouse (cKO, myeloid lineage), we previously found that A2B1 is required to induce inflammatory mediators during endotoxin shock and *Salmonella* infection (29). Mice lacking A2B1 exhibited higher levels of CFUs in peripheral immune organs and had reduced survival compared to WT controls (29). Similar phenotypes were reported for A2B1 cKO mice infected with enterohemorrhagic *E. coli,* and *Listeria monocytogenes* (54). Despite these compelling findings, the role of A2B1 in cell-intrinsic innate immunity remains unexplored.

Here, we integrate transcriptomic and molecular analyses to implicate A2B1 in host cell splicing changes during Mtb infection. We show that A2B1 plays a particularly critical role in controlling AS of the immunity-related GTPase Irgm1. We report that only the long isoform of Irgm1 promotes restriction of Mtb in macrophages and that the balance of the two *Irgm1* isoforms is differentially regulated in response to diverse inflammatory cues. These findings underscore the importance of AS in shaping antimycobacterial immunity and motivate further studies of functionally distinct roles of AS isoforms at the host-pathogen interface.

## RESULTS

### Mtb induces alternative splicing of functionally diverse genes during macrophage infec-tion

To date, alternative splicing (AS) has been implicated in the macrophage immune response to bacterial (*Salmonella enterica*, *Listeria monocytogenes*, *Mycobacterium tuberculosis* (Mtb)) and viral (DENV, HIV, Zika) pathogens (36, 55). Because macrophages serve as a replicative niche for Mtb and because AS transcripts have been identified has potential biomarkers for tuberculosis disease states in humans (35), we set out to refine our understanding of AS during Mtb infection of macrophages. Briefly, we infected bone marrow derived macrophages (BMDMs) isolated from C57BL/6 mice with Mtb (Erdman, MOI=5), isolated total RNA at 8h post infection, a time point at which pattern recognition receptors and Mtb-responsive signaling cascades are fully engaged but few infected cells have initiated programmed cell death pathways. We performed bulk RNA-seq (see Methods) and profiled AS changes alongside uninfected controls, using the annotation-agnostic MAJIQ-VOILA framework (56). Significant local splice variations (LSVs) were defined by a dPSI > 0.15 at a probability cutoff of 0.9 and a baseMean locus coverage > 50. Given our interest in how splicing impacts immune protein expression, we applied a protein-coding gate to our pipeline, where any gene of interest needed to be able to produce at least one protein-coding isoform as determined by our ORF scanner and CPAT (coding probability >1). To our knowledge, this represents one of the first reports of Mtb-induced AS in primary macrophages or in murine macrophages, providing an important foundation to explore the role of splicing in pre-clinical models of tuberculosis.

Of the 12,436 genes defined in the GRCm39 genome, MAJIQ-VOILA identified significant LSVs in ∼5% of genes (658 protein-coding genes of 688 detected), spanning the four canonical AS event types: alternative 5’ splice site (A5SS), alternative 3’ splice site (A3SS), exon skipping (ES), and intron retention (IR) (**Fig. 1A, Table S1**). ES (40.2%) and IR (38.0%) were preferred LSVs, with fewer A5SS (11.5%) and A3SS (10.3%) events detected (**Fig. S1A, Table S2**). Many genes had multiple AS changes—an average of 2.8 LSVs/gene (1,865 LSVs across 658 genes). Although many genes were transcriptionally up- or down-regulated during Mtb infection (see Methods), most AS-positive genes were not differentially expressed. Of the total 658 AS genes identified, 517 (4.16% of total expressed genes) underwent AS without differential expression, 65 (0.52%) underwent AS with decreased expression, and 76 (0.61%) underwent AS with increased expression (**Fig. 1A, Table S1**). The proportion of LSV type (ES, IR, A5SS, A3SS) was similar between differential expression-status groups (**Fig. 1B**). Together, these data suggest that although some transcriptionally induced innate immune genes are subject to AS, macrophages primarily use AS to post-transcriptionally regulate constitutively expressed genes in re-sponse Mtb infection.

**Figure 1.**
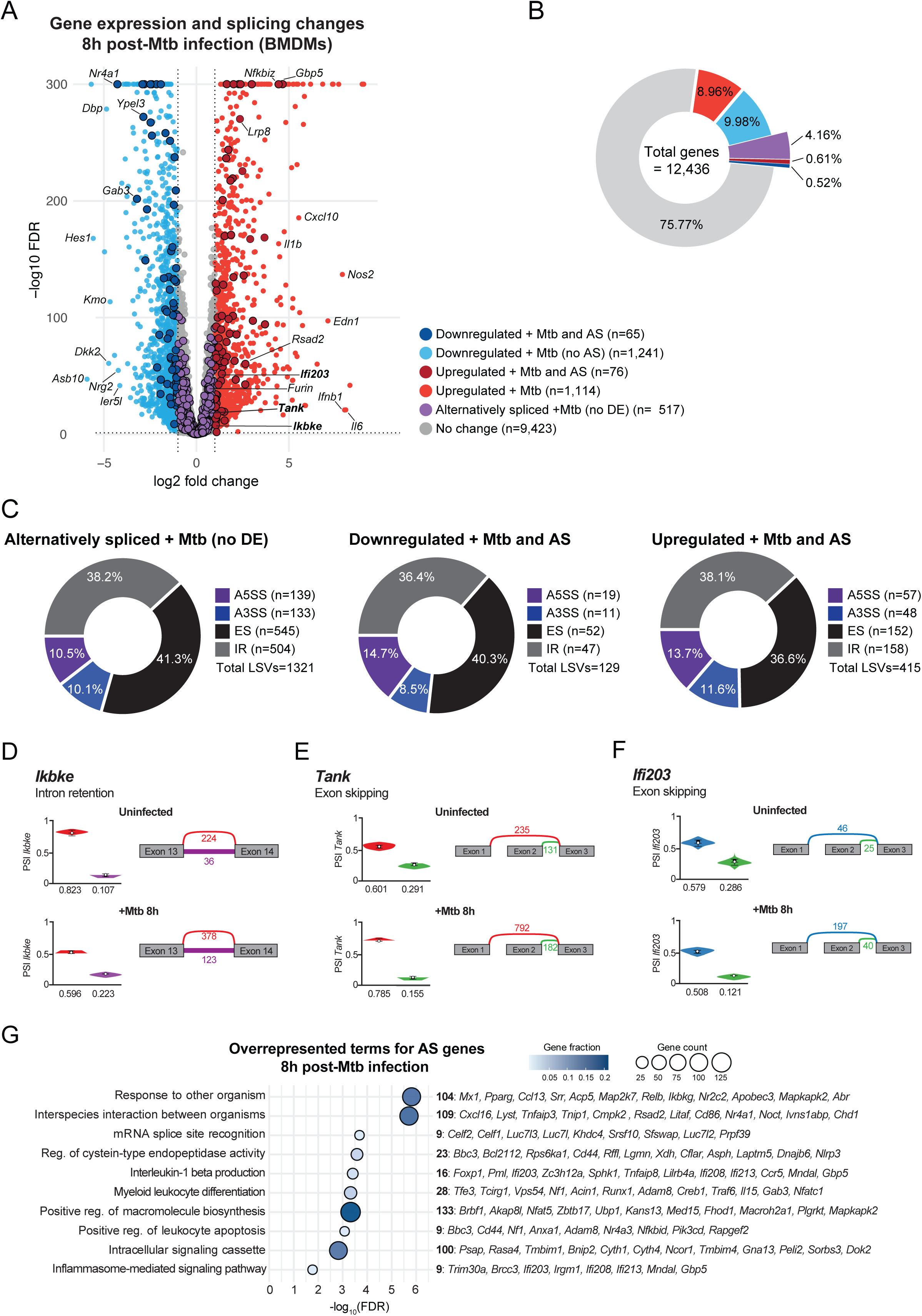
Mtb induces alternative splicing of functionally diverse genes during macro-phage infection. (A) Volcano plot of differentially expressed genes 8 hours post Mtb infection in WT Bone Marrow Derived Macrophages, genes with at least one significant (dPSI > 0.15, confidence threshold > 0.90) AS event are darkened. (B) Pie charts of distribution of categories of DEG and AS. (C) Pie charts of distribution of categories of significant Local Splice Variations for each DEG category. (D) MAJIQ PSI quantification and VOILA visualization of *Ikbke* junctions in uninfected (top) and Mtb-infected (bottom) BMDMs (E) As in D but for *Tank* (F) As in D but for *Ifi203* (G) Over Representation Analysis of pathways enriched for AS genes between uninfected and Mtb-infected WT BMDMs.

To identify genes that undergo the most dramatic shifts in AS during Mtb infection, we manually annotated genes with dPSI > 30% in each LSV category. Canonical immune genes meeting this threshold include the antiviral GTPase *Mx1*, the interferon-inducible guanylate binding protein *Gbp5*, and the IL-15 receptor subunit *Il15ra* (**Fig. S1B**). Notable immune genes with evidence of Mtb-regulated exon skipping included *Ikbke* (non-canonical kinase of IRF3 (57)), *Tank* (scaffold protein involved in TBK1-IRF3 signaling axis (58)), and *Ifi203* (non-canonical STING-dependent dsDNA sensor (59)) (**Fig. 1D-F**). To unbiasedly identify functional pathways enriched for genes that undergo AS during Mtb infection, we used Over Representation Analysis (ORA) referencing the GO: Biological Process term list, (**Fig. 1G**). Among the top 15 terms (FDR < 0.05, minimum pathway overlap of 2 genes, dPSI > 15% in any LSV category), nine were immune-related (e.g. *Response to other organism*, *Myeloid leukocyte differentiation*, and *Regulation of AIM2 inflammasome assembly*). Several of these pathways line up with prior studies of Mtb-infected THP1-1 cells that identified AS in genes related to autophagy (*Rab8b, Atg13. Response to other organism*) and inflammasome assembly (*Il1b. Regulation of AIM2 inflammasome assembly*) (34). We also detected AS of the inflammasome sensor *Nlrp3*, whose activity has been shown to be regulated by AS but has yet to be reported as AS in the context of Mtb infection (60, 61). Consistent with AS-mediated specialization of housekeeping genes during macrophage activation, we observed AS enrichment in genes involved in basic cellular and homeostatic functions—such as macromolecule biosynthesis, mRNA splice-site recognition, and developmental regulation (**Fig. 1G**). Enrichment profiles of AS genes and upregulated DEGs showed some overlap (**Fig. S1C-D**), hinting at dual regulation of pathways like *Intracellular signaling* and *Interspecies interaction between organisms* by AS and transcription. Collectively, these analyses reveal that Mtb infection of macrophages triggers a widespread, coordinated program of alternative splic-ing—driven largely by ES and intron retention (IR)—that is largely uncoupled from changes to total transcript abundance.

### The splicing factor hnRNPA2B1 regulates expression of inflammatory and type I interferon genes during Mtb infection

Our finding that distinct AS isoforms are generated in macrophages +/- Mtb argues that the process of splicing—and the factors that control splicing—are regulated during infection. RNA binding proteins, particularly those in the SR and hnRNP families, play key roles in dictating splice site choice and controlling toggling between AS isoforms (62). There is increasing appreciation that splicing factors themselves are functionalized during macrophage activation to generate specific isoforms that promote antimicrobial defenses (62). Based on our previous studies of the splicing factor hnRNPA2B1 in *Salmonella enterica*-infected mice, we hypothesized that A2B1 could play a role in the macrophage response to Mtb. To begin to test this, we harvested BMDMs from WT (LysM-cre/cre; hnRNPA2B1-+/+) and KO (LysM-cre/cre; hnRNPA2B1 fl/fl) mice. Loss of A2B1 expression at the protein and RNA levels was confirmed via immunoblot and RT-qPCR, respectively (**Fig. 2A**). WT and A2B1 KO BMDMs were infected with Mtb (MOI=5, 8 h) and RNA was collected for RNA-seq as in **Fig. 1** (see Methods). Differential expression analysis revealed 78 genes upregulated and 135 downregulated in A2B1 KO cells relative to WT during Mtb infection (**Fig. 2B**). To gain insight into potential functional outcomes of these gene expression changes, we performed GSEA using the HALLMARK database to identify pathways enriched for upregulated genes (**Fig. 2C**) or downregulated genes (**Fig. 2D**) in A2B1 KO BMDMs compared to WT. We observed widespread dampening of inflammatory pathways in Mtb-infected A2B1 KOs BMDMs (e.g. *TNF signaling*, *IL6 signaling*, and *Inflammatory response*) (**Fig. 2C, S2A-C**), alongside upregulation of genes in pathways related to mitochondrial homeostasis, reactive oxygen species, and type I interferon expression (**Fig. 2D, S2D-F**). Consistent with previous reports linking A2B1 to cell proliferation and cancer (43–46, 63), we also observed upregulation of genes related to the cell cycle, *Ccnb1*, and *Cdc20* in Mtb-infected A2B1 KOs relative to controls (**Fig. S2G**). Heatmaps of top DEGs in selected HALLMARK categories highlight the genes driving these terms: Inflammatory Response/TNF Signaling (*Il1b, Ptgs2, Nr4a3*), G2M Checkpoint (*Cdkn3, Aurka*), and Interferon Alpha Response (*Cmpk2, Rsad2, Cxcl10*) (**Fig. 2E**). RT-qPCR confirmed reduced inflammatory cytokine transcript induction and increased interferon and cell-cycle gene expression in A2B1 KO BMDMs at 8h post-Mtb infection (**Fig. 2F**). However, looking at a longer time-course of infection, transcriptional differences between genotypes appear to be transient: differences between A2B1-KO and WT macrophages are evident at 4h or 8h, but not 24h post-infection (**Fig. 2F**). Together, these findings suggest that hnRNPA2B1 plays a role in balancing the early induction of inflammatory and interferon-stimulated genes in response to Mtb.

**Figure 2.**
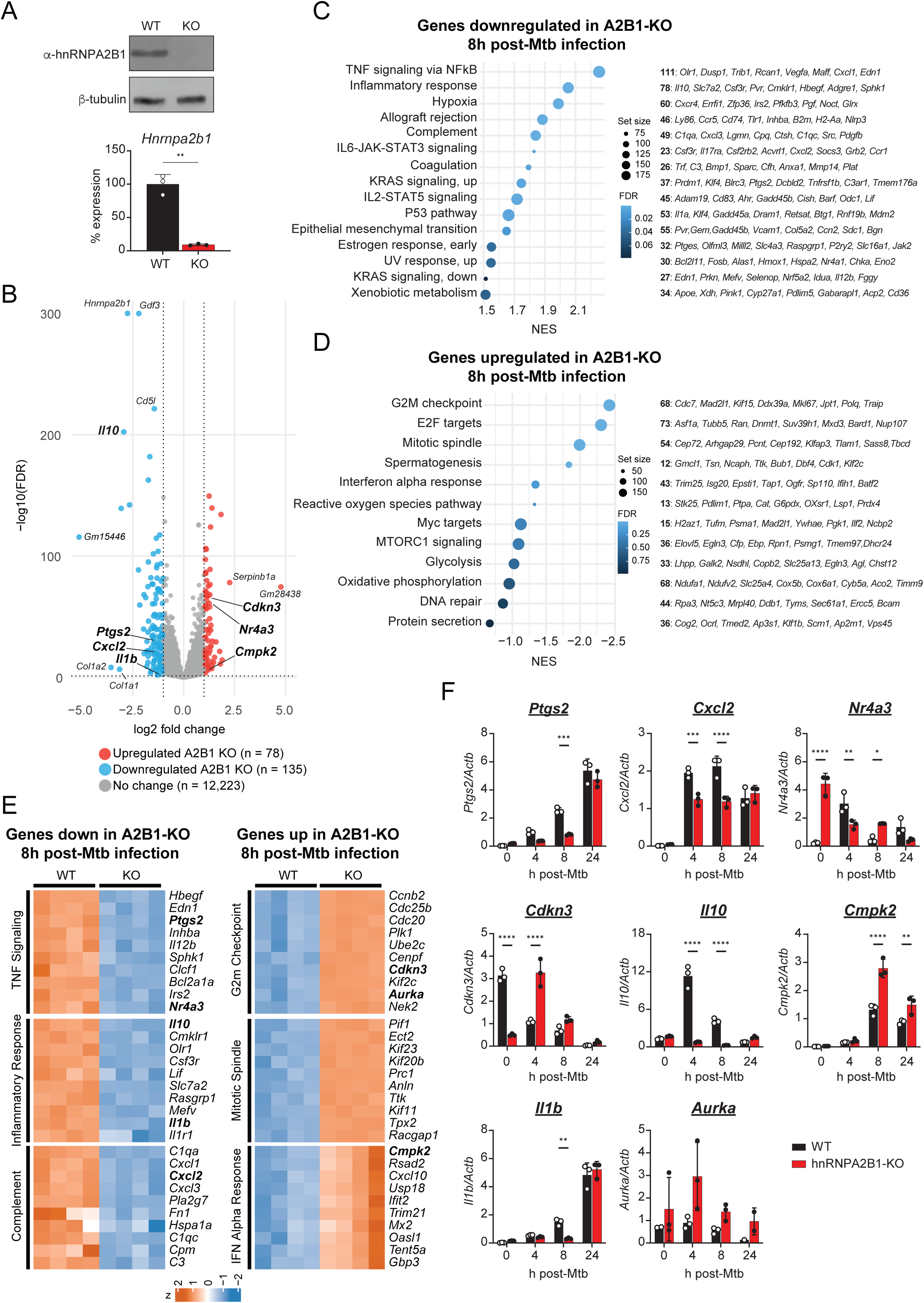
The splicing factor hnRNPA2B1 regulates expression of inflammatory and type I interferon genes during Mtb infection. (A) Immunoblot and RT-qPCR of hnRNPA2B1 in WT and hnRNPA2B1-KO BMDMs. (B) Volcano plot of differentially expressed genes (DEGs) between WT and hnRNPA2B1-KO BMDMs 8h post Mtb infection. (C) Gene Set Enrichment Analysis (GSEA) of pathways downregulated in DEGs in hnRNPA2B1-KO BMDMs during Mtb infection. (D) GSEA of pathways upregulated in DEGs in hnRNPA2B1-KO BMDMs during Mtb infection. (E) Heatmap of DEGs grouped by top HALLMARK terms in WT and hnRNPA2B1-KO BMDMs during Mtb infection. (F) RT-qPCR of selected DEGs in WT and hnRNPA2B1-KO BMDMs 0-,4-,8-, and 24- hours post Mtb infection. n=3.

### The splicing factor hnRNPA2B1 controls alternative splicing of genes during Mtb infection

Next, we asked how hnRNPA2B1 impacts AS during Mtb infection (8h). We identified 150 genes with significant AS changes (150 protein coding genes, of 151 AS detected), of which 144 showed no corresponding change in gene expression (**Fig. 3A, Table S3**). ORA of these 150 A2B1-dependent AS genes uncovered that seven of the top 15 enriched pathways are related to the macrophage immune response (*Regulation of the inflammasome, PRR signaling, Autophagosome maturation*) (**Fig. 3B**).

**Figure 3.**
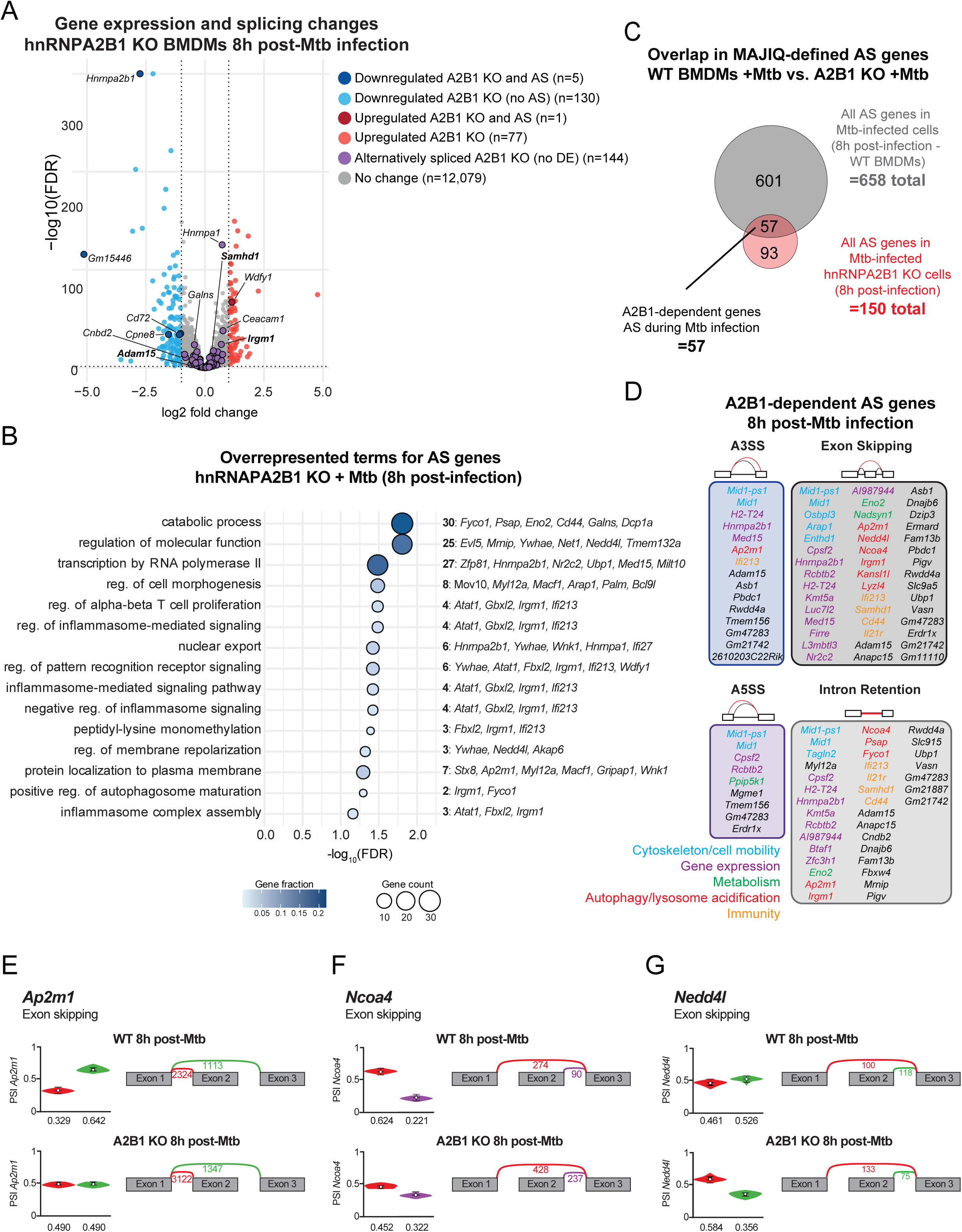
The splicing factor hnRNPA2B1 controls host alternative splicing decisions during Mtb infection. (A) Volcano plot of differentially expressed genes between WT and hnRNPA2B1-KO BMDMs 8h post Mtb infection, genes with at least one significant (dPSI > 0.15, confidence threshold > 0.90) AS event are darkened. (B) Over Representation Analysis of pathways enriched for AS genes between WT and hnRNPA2B1-KO BMDMs 8h post Mtb infection. (C) Overlap of significant AS genes in the WT uninfected vs 8h Mtb and 8h Mtb WT vs hnRNPA2B1-KO sets. (D) Lists of significant AS genes per LSV category from overlap region in C. (E) MAJIQ PSI quantification and VOILA visualization of *Ap2m1* in WT (top) and A2B1 KO (bottom) Mtb-infected macrophages (F) As in E but for *Ncoa4* (G) As in E but for *Nedd4l*

Comparison of A2B1-dependent AS events with those induced by Mtb in WT cells revealed 57 overlapping genes, representing Mtb infection-driven, hnRNPA2B1-dependent AS events. Most shared events were categorized as exon inclusion, consistent with the known role of A2B1 in promoting exon skipping (**Fig. 3C**). These hnRNPA2B1-dependent AS events occurred in genes involved in a variety of pathways, several with known links to antimycobacterial immunity (e.g. metabolism, autophagy, and lysosome acidification) (**Fig. 3D**). Notably we detected A2B1-dependent exon skipping in *Ap2m1* (**Fig. 3E**) *Ncoa4* (**Fig. 3F**) and *Nedd4l* (**Fig. 3G**).

*Ap2m1* encodes a component of the AP-2 clathrin adaptor complex that can be hijacked by *Listeria monocytogenes* to promote infection (64). *Ncoa4* is an E3 ubiquitin ligase involved in autophagy of ferritin-iron complexes (65). *Nedd4l* is an E3 ubiquitin ligase involved in autophagy, regulation of inflammation (66) and control of *Mycobacterium bovis* BCG replication (67). Collectively, these findings identify hnRNPA2B1 as a regulator of splicing decisions during Mtb infection, notably influencing isoform selection of genes acting in pathways with known links to mycobacterial immunity.

### Irgm1 has two isoforms that are differentially regulated upon immune activation of macrophages

Among hnRNPA2B1-regulated AS genes, *Irgm1* emerged as a compelling hit due to its essential roles in autophagy and antimycobacterial immunity (**Fig. 4A**). *Irgm1* (Immunity-Related GTPase family M protein 1) is a murine GTPase essential for host defense, with established roles in mitophagy, autophagy, and resistance to Mtb (68) (69, 70). *Irgm1* is functionally orthologous to human *IRGM*, which shares the IRG-like GTPase domain (PS51716) (**Fig. 4B-C, S3A**)*. Irgm1^-/-^* mice have been repeatedly shown to be incredibly sensitive to Mtb infection (69, 71, 72). While *in vivo* phenotypes have recently been attributed to misregulation of the type I IFN response (72), many foundational studies implicate Irgm1 in cell-intrinsic control of mycobacteria via autophagy (71–74). Relevant to our studies of AS in BMDMs, the murine *Irgm1* locus produces two protein-coding isoforms via exon 2 inclusion: Irgm1-long, which contains a 16-aa N-terminal extension, and Irgm1-short, which lacks this region. Prior work suggests that both isoforms can localize to the Golgi but only Irgm1-long localizes to lysosomes (74). Beyond altered subcellular localization, no functional differences have been attributed to the two Irgm1 isoforms.

**Figure 4.**
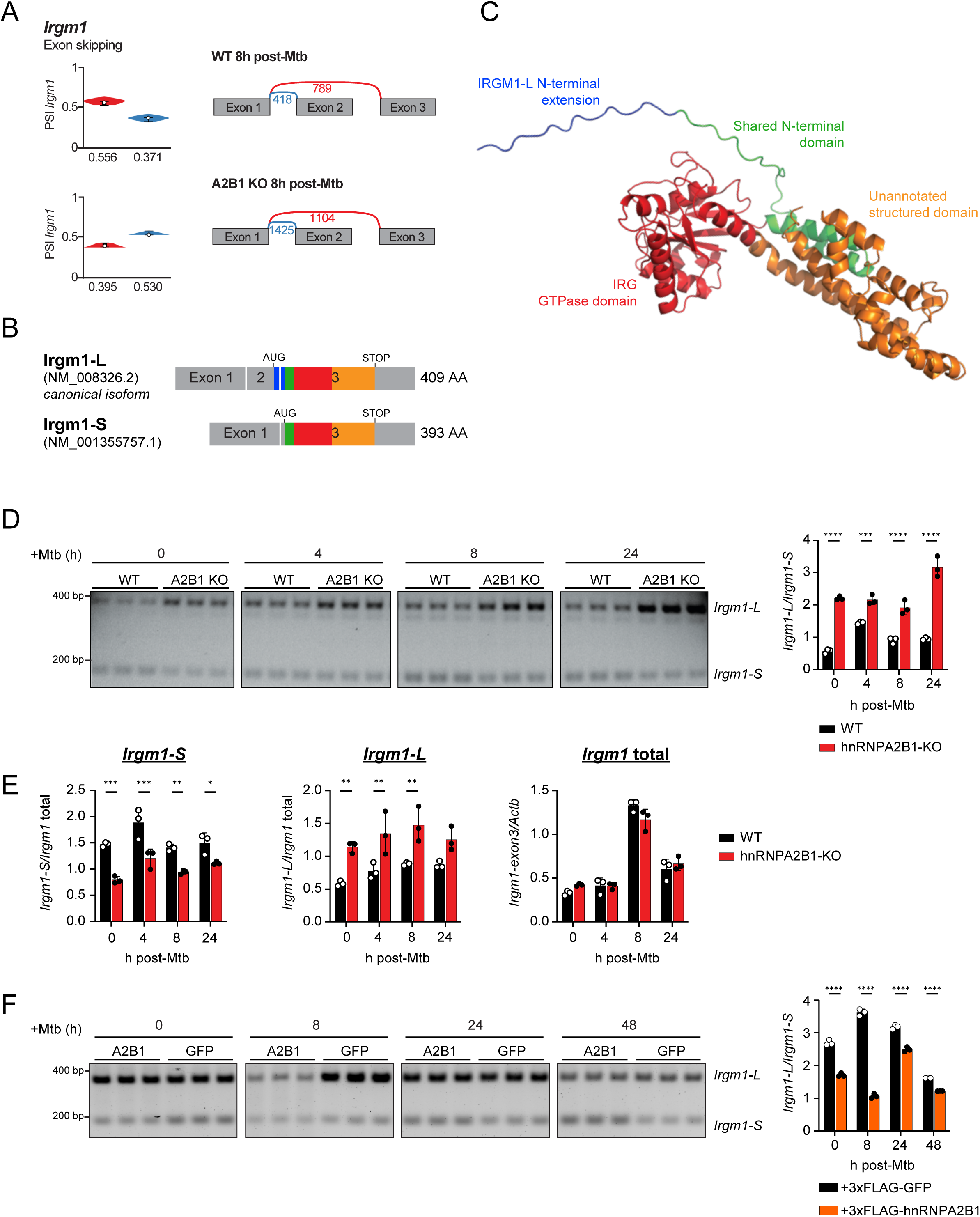
Irgm1 is an anti-mycobacterial, autophagy protein and is alternatively spliced in hnRNPA2B1-KO BMDMs (A) MAJIQ PSI quantification and VOILA visualization of *Irgm1* in WT (top) and A2B1 KO (bottom) Mtb-infected macrophages (B) Diagram of Irgm1 isoform exon usage. (C) Predicted Alphafold structure of Irgm1 with annotated regions highlighted. (D) Semiquantitative RT-PCR of *Irgm1* exon 2 inclusion in WT and hnRNPA2B1-KO BMDMs 0-, 4-, 8-, and 24-hours post Mtb infection. Each sample in biological triplicate. Quantification represented as band intensity of *Irgm1-long* (top) over band intensity of *Irgm1-short* (bottom) shown on the right. n=3. (E) RTqPCR of *Irgm1* isoforms and total transcript abundance in WT and hnRNPA2B1-KO BMDMs 0-, 4-, 8-, and 24- hours post Mtb infection. n=3. (F) Semiquantitative RT-PCR of *Irgm1* exon 2 inclusion in GFP and hnRNPA2B1 overexpression iBMDMs 0-, 8-, 24-, and 48-hours post Mtb infection. Quantification represented as band intensity of *Irgm1-long* (top) over band intensity of *Irgm1-short* (bottom) shown on the right. n=3.

To validate our RNA-seq findings and examine the dynamics of Irgm1 AS, we used semi-quantitative RT-PCR to quantify the long isoform (contains exon 2; generates a 394 bp PCR product) and the short isoform (exon 2 is skipped; generates a 175 bp PCR product) in RNA isolated from WT and A2B1-KO BMDMs at 4, 8, and 24h post-Mtb infection (**Fig. S3B)**. Consistent with our RNA-seq data, A2B1 KO macrophages accumulated the long isoform in all timepoints relative to controls (**Fig. 4D**). RT-qPCR confirmed these results (**Fig. 4E, S3C**). Notably, total *Irgm1* transcript abundance remained unchanged between genotypes, indicating that A2B1 controls splicing rather than transcriptional regulation of this gene (**Fig. 4E**). Conversely, A2B1 overexpression (specifically of the A2 isoform, the dominant form of the protein expressed in macrophages) (**Fig. S3D**) shifted isoform usage toward *Irgm1-short*, reinforcing its role in exon 2 exclusion (**Fig. 4F**). Immunoblot demonstrated increased Irgm protein in hnRNPA2B1-KO cells during infection, although we are unable to distinguish Irgm1 isoforms with this antibody, both due to the small size difference between the two protein isoforms and the fact that the antibody also recognizes Irgm2 and Irgm3 (**Fig. S3E**). Collectively, these findings establish hnRNPA2B1 as a key regulator of *Irgm1* splicing in macrophages.

Previous reports have shown that Irgm1-long and -short differ in their subcellular localization (74). To further explore this, we fed the sequences of Irgm1-long and -short into DeepLoc2.1, a software that predicts the subcellular localization of proteins (75). Although none of the localization predictions meet the program’s significance threshold, DeepLoc2.1 predicts that Irgm1-short is most likely found in the cytosol and plasma membrane whereas Irgm1-long is most likely targeted to lysosomes (**Fig. S3F**). To test these predictions, we transfected N-terminal FLAG tag Irgm1-long, Irgm1-short, or a GFP overexpression control into HEK293T cells (**Fig. S3G**) and monitored subcellular localization of each construct via immunofluorescence microscopy. We found that anti-FLAG staining of 3xFLAG-Irgm1-short neatly outlines the cell membrane of the cell and aggregates in small puncta throughout the cytosol (**Fig. S3H**). 3xFLAG-Irgm1-long, on the other hand, aggregates in small puncta but also exhibits a strong perinuclear enrichment (**Fig. S3H**). We did not observe any striking colocalization between either isoform with specific cellular organelles (data not shown). Overall, staining of both isoforms was consistent with previous reports of IRGM marking a variety of cellular compartments (e.g. Golgi, ER, phagosomes, endosomes) as “self,” for protection against GKS-protein mediated damage (74, 76–79).

### Overexpression of *Irgm1-short* inhibits Mtb restriction in macrophages

To understand the functional consequences of *Irgm1* AS in the context of antimycobacterial immunity, we generated WT iBMDM cell lines that stably overexpress each isoform (**Fig. 5A**) and infected them with Mtb-lux, monitoring infectious burden via luciferase assay over 5 days (as in (80)). Relative to the GFP control, cells overexpressing Irgm1-long had lower bacillary burdens whereas cells overexpressing Irgm1-short had higher burdens (**Fig. 5B**). To rule out the possibility that differences in Mtb-lux signal could be due to altered sensitivity of the two isoform-expressing cell lines to undergo cell death, we monitored propidium iodine incorporation over a 24h time-course of Mtb infection. Cell death kinetics were similar in cells expressing GFP, Irgm1-long and Irgm1-short (**Fig. S4A**). Likewise, monolayers remained intact over the entire 5-day time-course (**Fig. S4B**), arguing that Irgm1-long restricts Mtb replication via a cell death-independent mechanism.

**Figure 5.**
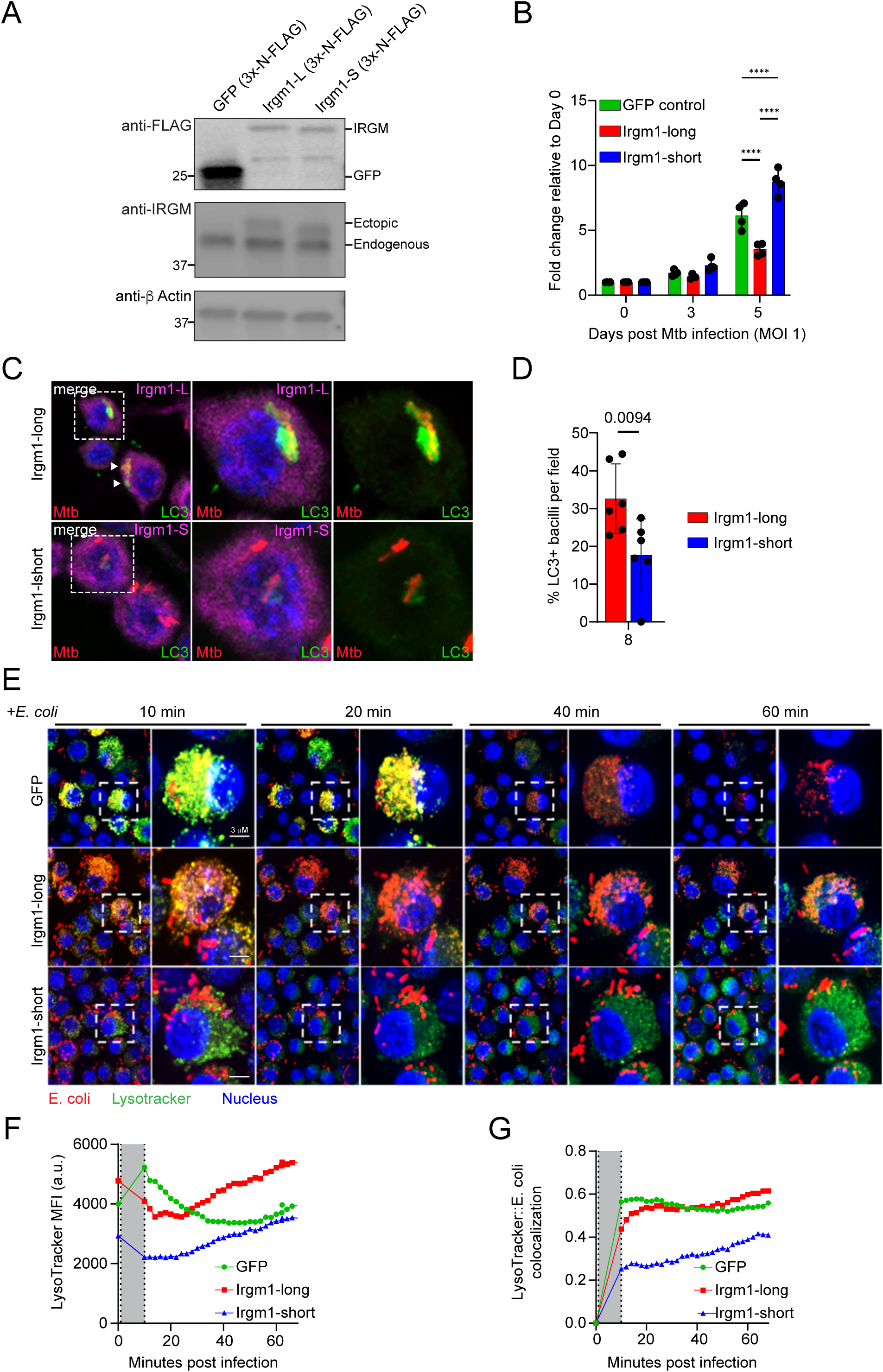
Irgm1 isoforms have different capacity for restricting Mtb replication in iBMDMs (A) Immunoblot of IRGM in GFP, FLAG-Irgm1-long, and FLAG-Irgm1-short stably expressed in WT iBMDMs. (B) Fold replication of Mtb in GFP, Irgm1-long, or Irgm1-short stable expression iBMDMs 0-, 3-, and 5- days post infection, measuring luminescence generated via Mtb-lux ABCDE. Shown as fold change relative to day 0. n=4. (C) Immunofluorescence microscopy of Irgm1-isoforms (FLAG, pink), LC3 (green), and Mtb (mCherry, red) 8 hours post-Mtb infection in Irgm1-long and Irgm1-short-expressing iBMDMs. (D) Quantification of (C) (E) Live-cell fluorescence microscopy of *E. coli* (mCherry, red), LysoTracker (green), and nucleus (blue) 0 to 68 minutes post *E. coli* infection in GFP, Irgm1-long, and Irgm1-short iBMDMs. (F) Quantification of (E); total LysoTracker per field at baseline and through duration of infection. Grey bar represents infection step (10 minutes). (G) Quantification of (E); Pearson Colocalization Coefficient of LysoTracker with mCherry at baseline and through duration of infection. Grey bar represents infection step (10 minutes).

Given that the two isoforms differ in their ability to control Mtb survival/replication (**Fig. 5B**), we reasoned that Irgm1-short isoform expressing cell lines may be defective in one or more steps of selective autophagy, a process known to restrict Mtb replication (81) and linked to IRGM1 (72, 79, 82). To begin to test this, we used immunofluorescence microscopy to quantify abundance and localization of autophagy factors at 8 hours post-infection of Irgm1 isoform-expressing iBMDMs with mCherry Mtb. Consistent with Mtb-lux results in **Fig. 5B**, we measured increased bacilli area in Irgm1-short iBMDMs at 24 hours post-infection by quantifying mean fluorescence intensity measured across segmented bacillus area (**Fig. S4C**). We have previously shown that LC3 is recruited to a population of Mtb-containing phagosomes at 6-8h post-infection and is degraded during lysosomal turnover of autophagosomal cargo (81). Early studies of IRGM1 from the Deretic lab demonstrated fewer LC3+ mycobacteria in Irgm1 knockdown macrophages (83). Consistent with this finding, we measured higher LC3 MFI per cell and higher percentage of LC3+ Mtb bacilli in Irgm1-long expressing cells relative to Irgm1-short at 8h post infection (**Fig. 5C-D, S4D**). Because LC3+ membranes are turned over when autophagosomes fuse with lysosomes, fewer LC3+ bacilli in Irgm1-short iBMDMs could represent either impaired autophagosomal targeting or enhanced lysosomal degradation. To begin to distinguish these models, we quantified Mtb colocalization with the lysosomal marker, LAMP1. Although total LAMP1 signal was the same in Irgm1-long vs. -short expressing iBMDMs (**Fig. S4E**), Irgm1-short cell showed significantly less LAMP1-Mtb colocalization 8h post-infection (**Fig. S4F**), supporting a model whereby expression of Irgm1-short impairs LC3+ tagging of Mtb, thereby reducing lysosomal targeting of autophagosomes rather than accelerating it.

Given that Irgm1-long is reported to colocalize with lysosomes (74), we wanted to test if these isoforms could differentially impact lysosomal function in macrophages. Because our BSL3 lacks live-cell imaging capabilities, we turned to another intracellular bacterial model. Briefly, we infected Irgm1-long, Irgm1-short, and GFP-expressing iBMDMs with a lab strain of mCherry-expressing *E. coli* (DH5a) and monitored acidification of *E. coli*-containing compartments via live-cell imaging of LysoTracker. Because non-pathogenic E. coli has not evolved any strategies to avoid delivery to lysosomes, macrophages rapidly target bacili to lysosomes for destruction. We found that cells expressing Irgm1-long had a higher lysotracker MFI at baseline (**Fig. S4G-H**) and over the duration of the infection (**Fig. 5E, F**) as well as higher colocalization of lysotracker with the mCherry *E. coli* (**Fig. 5E, G**) compared to cells expressing Irgm1-short (see also **Video S1**). These findings suggest that Irgm1-short is sufficient to impair lysosomal function, both maintenance of low pH compartments and fusion of lysosomes with bacterial-containing endosomes.

### Pathogen-derived and inflammatory stimuli differentially regulate *Irgm1* AS

Upon phagocytosis, Mtb engages the host macrophage through PRRs (TLR1/2/6, cGAS-STING) and through release of virulence factors (via various secretion systems). After PRR engagement, infected macrophages release and respond to various cytokines, notably IFN-β, IFN-γ, TNF-α.

Given the functional consequences of the splicing decision of *Irgm1*, we set out to characterize how Mtb-relevant immune stimuli influence *Irgm1* splicing decisions. Using the same semi-quantitative RT-PCR approach employed in **Fig. 4D**, we first asked whether upregulation of *Irgm1-long* was dependent on Mtb’s ESX-1 secretion system, which controls cytosolic access and delivery of key virulence factors (84, 85). Compared to WT Mtb-infected BMDMs, which express high amounts of *Irgm1-long* relative to -short at 8 and 24h post-infection (**Fig. 6A**), ΔESX-1 Mtb infected macrophages make more of the short isoform (**Fig. 6B-C**), suggesting that Mtb secreted factors and/or cytosolic access of Mtb promotes generation of *Irgm1-long* during infection. Treatment of Mtb-infected cells with the STING inhibitor H-151 had no effect on Irgm1 AS until very late timepoints (48h post-infection), arguing against early upregulation of Irgm1-long being dependent on cGAS-STING signaling (**Fig. S5A, D, E**).

**Figure 6.**
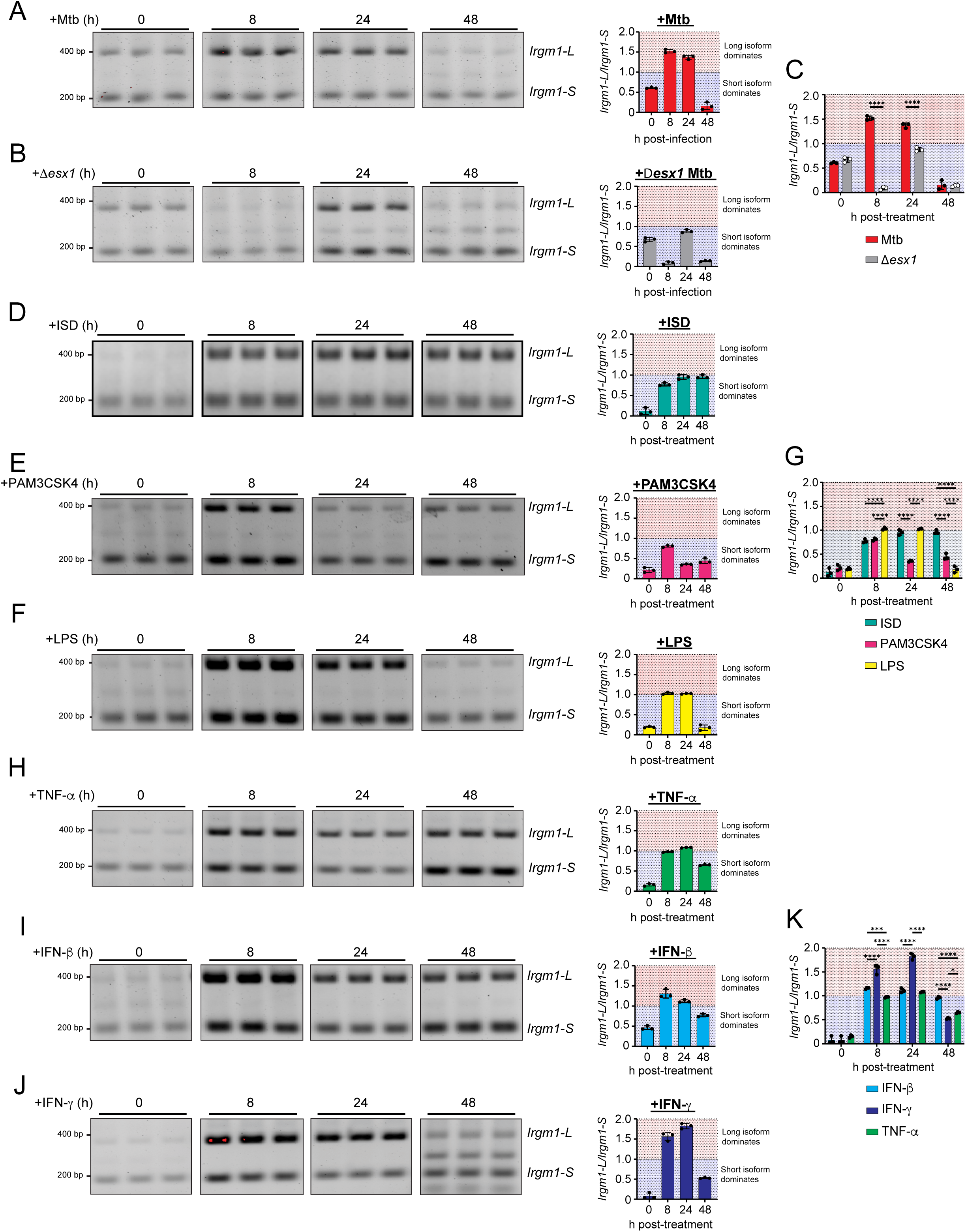
Irgm1 splicing in BMDMs is differentially responsive to distinct innate immune cues (A) Semiquantitative RT-PCR of *Irgm1* exon 2 inclusion in WT and hnRNPA2B1-KO BMDMs 0-, 8-, 24-, and 48-hours post Mtb infection (MOI = 5). Quantification on right represented as band intensity of *Irgm1-long* (top) over band intensity of *Irgm1-short* (bottom). n=3 (B) As in B, but 0-, 8-, 24-, and 48-hours post-infection with dESX1-Mtb (MOI = 5) (C) Side by side comparison of Irgm1-long to -short ratios between (A) and (B). (D) As in A, but post-transfection with ISD (1 ug/mL, 0-, 8-, 24-, 48- hours) (E) As in A, but post-stimulation with PAM3CSK4 (1 ug/mL, 0-, 8-, 24-, 48- hours) (F) As in A, but post-stimulation with LPS (10 ng/mL, 0-, 8-, 24-, 48- hours) (G) Side by side comparison of Irgm1 Irgm1-long to -short ratios between (D), (E), and (F). (H) As in B, but post-stimulation with TNF-α (20 ng/mL, 0-, 8-, 24-, 48- hours) (I) As in B, but post-stimulation with IFN-β (50 IU/mL, 0-, 8-, 24-, 48- hours) (J) As in B, but post-stimulation with IFN-γ (50 IU/mL, 0-, 8-, 24-, 48- hours) (K) Side by side comparison of Irgm1 Irgm1-long to -short ratios between (H), (I), and (J).

We next assessed how individual activation of Mtb-relevant PRRs influences *Irgm1* splicing. Briefly, we transfected BMDMs with interferon stimulatory DNA (ISD) (cGAS agonist, [1 ug/mL]) or stimulated BMDMs with PAM3CSK4 (TLR1/2/6 agonist, [1 ug/mL]). Consistent with inhibiting STING during Mtb infection, we found that ISD promoted roughly equal amounts of *Irgm1-long* and *Irgm1-short* (**Fig. 6D**). We found that stimulation with PAM3CSK4 also promotes similar amounts of each isoform (**Fig. 6E**). Although not relevant during Mtb infection, we also asked how engagement of TLR4 via LPS altered *Irgm1* isoform abundance. Consistent with TLR4 and TLR2 signaling sharing many components, LPS (10 ng/ml) also promoted similar levels of *Irgm1-long* and *-short* (**Fig. 6F-G**).

Finally, we tested how different secreted cytokines, sensed in an autocrine or paracrine fashion by macrophages, impact *Irgm1* AS. TNF-α (20 ng/mL), an abundant and critical pro-inflammatory cytokine rapidly induced upon Mtb infection of macrophages (86), promoted similar amounts of *Irgm1-long* and *-short* (**Fig. 6H**). Interesting, although the anti-bacterial type II inter-feron IFN-γ (50 IU/mL) and the pro-bacterial type I IFN IFN-β (50 IU/mL) transcriptionally upregulate *Irgm1* (73, 87), only IFN-γ promoted generation of *Irgm1-long* relative to short (**Fig. 6I-J**). Interestingly, blocking IFNAR with a monoclonal antibody during Mtb infection was sufficient to push Irgm1 splicing towards the long isoform, suggesting that IFNAR signaling negatively regulates generation of Irgm1-long (**Fig. S5B, D, E**). Given that type I IFN has been repeatedly shown to promote Mtb pathogenesis in macrophages and *in vivo* (88–90), it is tempting to speculate that failure to generate Irgm1-long in a type I IFN-dominant cytokine milieu contributes to these phenotypes. Together, these data show that *Irgm1* AS can be differentially regulated in response to distinct inflammatory cues, and suggest that IFN-γ, a potent anti-bacterial cytokine, is unique in its ability to promote the Mtb-restrictive isoform of IRGM1. Because no single agonist treatment recapitulated the Mtb-induced dynamics of Irgm1 isoform expression, our data also argue that regulation of Irgm1 AS during Mtb infection likely involves multiple inputs.

## DISCUSSION

Tight control of antimicrobial proteins is required to preserve cellular homeostasis while maintaining the capacity to mount effective host defenses. The family of dynamin-like immunity-related GTPases (IRGs), of which Irgm1 is a member, is divided into two groups: GKS “effector proteins” and GMS “regulatory proteins.” The presence of GMS proteins like Irgm1 on intracellular membranes is thought to mark them as “self,” preventing the activation of GKS proteins and subsequent GKS-mediated damage (91, 92). To date, studies of Irgm1 in host defense and autoimmunity have largely focused on its canonical long isoform. Here, we demonstrate that AS plays an important role in toggling *Irgm1* between a long isoform and a short isoform that lacks an N-terminal domain. By implicating the splicing factor hnRNPA2B1 in balancing the abundance of Irgm1-long vs. short and showing that Irgm1-long abundance is enhanced by Mtb in-fection and IFN-γ treatment, this work advances our understanding of how macrophages use AS to regulate antimycobacterial innate immunity.

Although differential subcellular localization of Irgm1-long and short has been previously reported (74), our data shed new light on how these proteins may function differently during bacterial infection of macrophages. Most notably, we show that expression of Irgm1-long promotes restriction of Mtb replication in macrophages, while expression of Irgm1-short creates a more permissive niche for Mtb (**Fig. 5B**). The mechanisms underlying these phenotypes appear to be somewhat complex. Unlike Irgm1-long, expression of Irgm1-short is not sufficient to promote LC3+ recruitment to Mtb bacilli (**Fig. 5C-D**). It is however, sufficient to inhibit lysosomal function, as evidenced by lack of colocalization between *E. coli* and the pH-sensitive dye LysoTracker (**Fig. 5E-G**). Lysosomal defects have previously been reported in mouse embryonic fibroblasts lacking Irgm1 (76). Such defects are attributed to mislocalization of GKS proteins (e.g. Irga6, Irgb6, Irgb10) to lysosomes and subsequent lysosomal dysfunction (evidenced by impaired dequenching of the pH-sensitive fluorescent dye DQ-BSA) (76). Although we cannot distinguish whether Irgm1-short inhibits lysosome acidification, fusion of lysosomes with internalized *E. coli*, or both, our data suggest that Irgm1-short expression generally phenocopies loss of the *Irgm1* gene locus. These findings support a model whereby Irgm1-short opposes the function of Irgm1-long, acting as an internal rheostat to tune lysosomal capacity in response to infection or other external cues. Given that AS impacts the inclusion of a N-terminal targeting region, yet leaves the GTPase domain intact, this differential subcellular localization may sequester Irgm1 binding partners away from compartments of the cell where they are needed to function in cellular processes like autophagy (**Fig. 7**).

**Figure 7.**
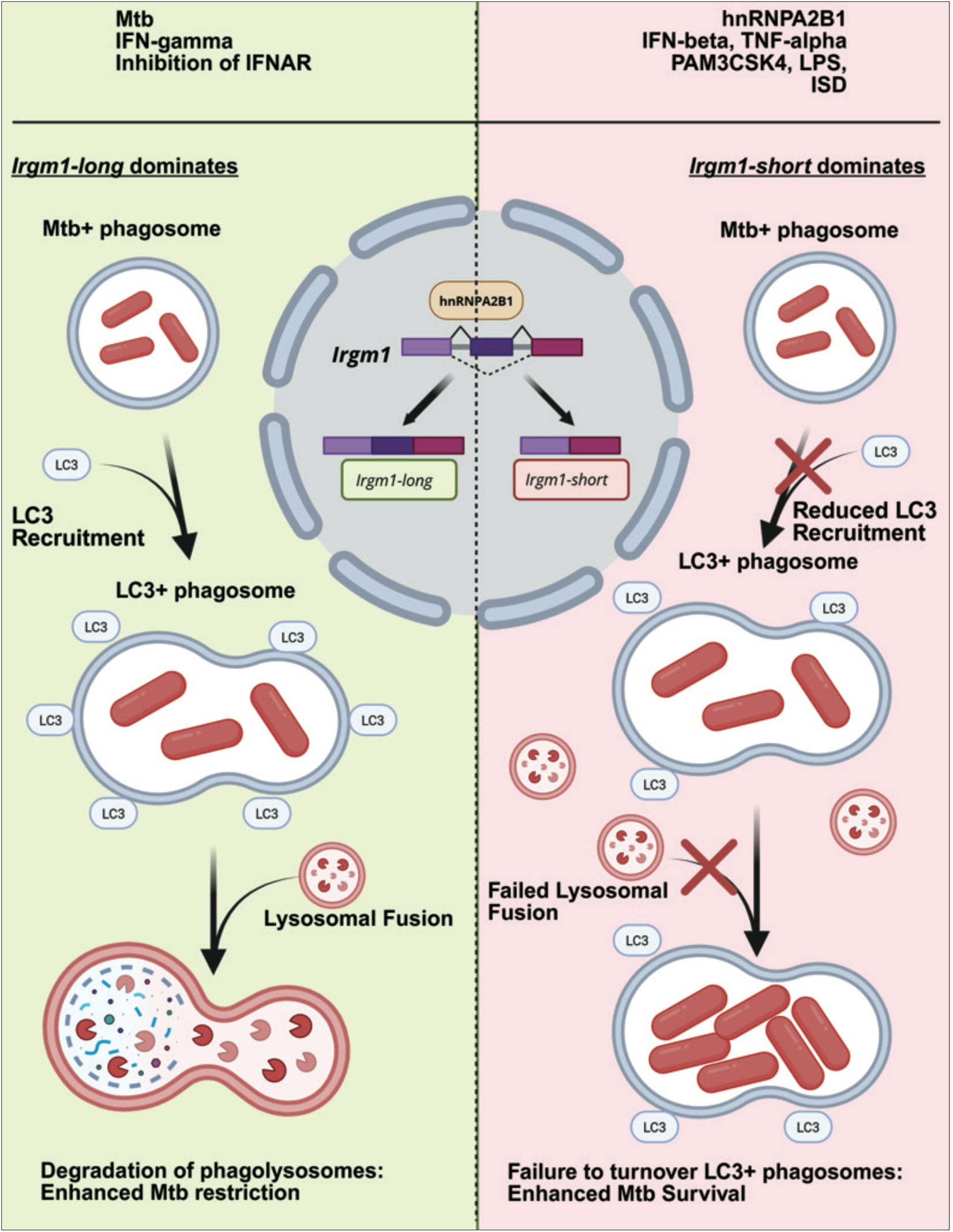
Model: Irgm1 isoforms differentially impact autophagic clearance of Mtb In stimulated macrophages, *Irgm1* transcripts increase in abundance and are alternatively spliced into long or short isoforms by hnRNPA2B1 (with the presence of A2B1 favoring skipping of exon 2 and generation of the short isoform). The decision to make Irgm1-short vs. Irgm1-long can be influenced by different immune stimuli, with Mtb infection, IFN-γ, and IFNAR blocking promoting Irgm1-long, and PRR stimulation (cGAS-STING, TLR1/2/6) and cytokine treatment (TNF-α, IFN-β) eliciting more balanced expression of Irgm1-long and short. Irgm1-long supports autophagy, enabling LC3 recruitment to Mtb-containing phagosomes and subsequent lysosomal fusion, thereby restricting bacterial replication. In contrast, Irgm1-short impairs both LC3 and lysosomal recruitment, leading to defective autophagy and failure to control intracellular Mtb. Created in BioRender. Chapman, M. (2026) https://BioRender.com/3ldv1ks

It is not entirely clear why we see more LC3+ Mtb bacilli in Irgm1-long vs. Irgm1-short cells. Although our finding is consistent with Irgm1 knockdown cells harboring fewer LC3+ BCG (83), early reports of Irgm1 recruitment to Mtb phagosomes (93) have largely been dismissed (74). We also do not see any evidence for Irgm1-long or -short colocalizing with Mtb (data not shown). We propose that instead of Irgm1 physically directing LC3+ to Mtb phagosomes, its loss/overexpression misregulates components of the autophagy machinery. The Deretic lab has shown that IRGM1 interacts with core components of the autophagy machinery and IRGM1 knockdown decreases steady state levels of pULK-1, ATG5, and ATG16L1 in U937 cells (94).

Overexpression of Irgm1-long, but not Irgm1-short, may increase the stability or availability of one or more of these factors, such that they are ready to participate in antimycobacterial selective autophagy. Altered lysosomal function may also influence the number of LC3+ bacilli in Irgm1-long vs. -short expressing cells, by misregulating autophagolysosome formation or turnover of targeted cargo.

Establishing hnRNPA2B1 as a key regulator of Irgm1 isoform selection represents a considerable advance in our understanding of how AS can direct antimycobacterial immunity. Our finding that *Irgm1* AS is regulated by distinct immune agonists hints at the potential for A2B1 itself to be regulated. Various stressors (e.g. heat shock, genotoxic stress) are known to alter the phosphorylation status and subcellular localization of SR/hnRNP family members to control AS decisions (95–98). Analogous regulation has been reported in macrophages responding to immune agonists and/or infection (99, 100). Our own data demonstrate that LPS-directed p38-dependent phosphorylation of hnRNP M at S574 alters its capacity to associate with the IL6 genomic locus and repress IL6 maturation (12). We propose that post-translational modification of A2B1, down-stream of specific signals, toggles AS of Irgm1. High-throughput proteomics studies have identified many PTM sites on hnRNPA2B1 (**Fig. S5F**)(101) and quantitative proteomics of Mtb-infected BMDMs hint at differential phosphorylation of A2B1 at Y244 (102). The outstanding question then, is what signals could direct this modification and others? We found that Mtb and IFN-γ selectively promote splicing of *Irgm1-long* (**Fig. 6A, J**). In the case of Mtb, this required an intact ESX-1 secretion system, suggesting cytosolic access, the detection of Mtb-derived PAMPs like dsDNA/dsRNA, and/or secreted virulence proteins are somehow integrated to signal A2B1 to make more Irgm1-long relative to -short. The fact that we were unable to recapitulate dominant expression of Irgm1-long with any single Mtb-derived agonist suggests that the host integrates multiple signals during Mtb infection to regulate A2B1 splicing of *Irgm1*. It is notable that the only cytokine (of those tested) that promoted the generation of Irgm1-long was IFN-γ, a well-established inducer of IRGM1-dependent antimycobacterial immunity. Despite sharing many signaling components, IFN-γ is seemingly better than IFN-β at promoting Irgm1-long, suggesting that signaling downstream of IFNGR somehow delivers a unique signal to A2B1. Molecular dissection of these signaling cascades via genetics and inhibitors will provide new insights into how macrophages differentially regulate AS to generate a specialized antibacterial proteome.

## METHODS

### Mice and ethics statement

C57BL/6J (Jackson Laboratories, stock no. 00064) and hnRNPA2B1-KO mice (a gift from the Carpenter laboratory, University of California, Santa Cruz) were bred and maintained under approved protocols of the Texas A&M College of Medicine and Vanderbilt University Medical Center’s Institutional Animal Care and Use Committees. Age- and sex-matched littermates (8–16 weeks) were used for all experiments. Mice were housed on a 12-h light/dark cycle with ad libitum access to food and water.

### Cell culture

Immortalized bone-marrow-derived macrophages (iBMDMs) and HEK293T cells were cultured in high-glucose DMEM (Thermo Fisher) supplemented with 10 % FBS (Millipore) and 0.2 % HEPES at 37 °C in 5 % CO₂. All cell lines tested negative for mycoplasma contamination.

### Generation of bone-marrow-derived macrophages

Primary BMDMs were differentiated from femurs and tibias of 8–12-week-old C57BL/6J mice as previously described (Jove et al.). Briefly, marrow progenitors were cultured for 6 days in DMEM containing 10 % heat-inactivated FBS and 20 % L929-conditioned M-CSF. Cells were harvested in PBS–EDTA, washed, and replated for infection or stimulation assays.

### Immortalization and stable expression lines

WT BMDMs were immortalized by J2 CRE retroviral transduction to generate iBMDMs (Salih et al., 2025). Stable RAW 264.7 and iBMDM lines expressing GFP-FLAG, Irgm1-short-FLAG, or Irgm1-long-FLAG were produced using pLenti PGK-Puro DEST vectors followed by puromycin selection. Expression was confirmed by anti-FLAG immunoblotting.

### Mtb infections

All pathogen work was conducted under appropriate biosafety conditions. *Mycobacterium tuberculosis* Erdman and LuxABCDE-Erdman strains were grown in Middlebrook 7H9 broth with OADC, Tween-80, and glycerol. BMDMs were infected at an MOI of 1, centrifuged (1000x RPM VWR MegaStar 4.0, 10 minutes) briefly to synchronize infection, and washed 2 h later. CFU counts were obtained by serial dilution on 7H10 agar; luminescence (RLU) was measured on a TECAN Spark reader.

### Innate immune stimulations

BMDMs or RAW cells were stimulated for 4 h with 100 ng/mL LPS (InvivoGen) or 1 ug/mL Pam3CSK4 (InvivoGen), or transfected with 1 µg/mL interferon-stimulating DNA (ISD) (Patrick Lab) or poly(I:C) (InvivoGen) using Lipofectamine. Cytokine stimulations used 50 IU/mL IFN-β (PBL Assay Science), 50 IU/mL IFN-γ (PBL Assay Science), or 20 ng/mL TNF-α (PeproTech). Inhibition of innate immune pathways was performed by incubating cells in 5 uM H-151 (STING) (Cayman) or 160 IU/mL MAR1-5A3 (eBioscience) (ThermoFisher) for 5 hours prior to infection, then leaving inhibitor in culture for remainder of infection.

### RNA isolation and sequencing

RNA was extracted in TRIzol and purified with Direct-zol RNA Miniprep kits (Zymo Research) including on-column DNase digestion. Library preparation and sequencing (paired-end 150 bp, NovaSeq 6000) were performed by the Baylor College of Medicine GARP Core. Reads were trimmed with fastp (v 1.0.1) (103) and aligned with STAR (v 2.7.11b) (104) to the GRCm39/Gen-code v44 reference. Differential expression was analyzed with featureCounts (v2.1.1) (105) and DESeq2 (v1.46.0)(106), and alternative splicing was evaluated using MAJIQ and VOILA (v 2.5.11; |ΔPSI| ≥ 0.15, P ≥ 0.9)(56, 107). Protein coding status was determined by IsoformSwitchAnalyzeR (v) (108) to screen isoforms for the longest ORF, CPAT (v 3.0.5)(109) was then used to remove ORFs with low probabilities of being translated (< 0.5), the gene locus was considered protein-coding if at least one isoform met these criteria. DeepLoc2.1 (75) was used to determine subcellular localization. Gene-ontology was performed using ClusterProfiler (110), ORAs referenced the GO: Biological Process database (111), GSEAs referenced the HALLMARK database0 (112, 113). Predicted structures for isoforms were obtained from the AlphaFold Protein Structure Database (114, 115), .pdb files were visualized for annotation using the Pymol software (v2.5.4) (116). Plots and statistical analyses were generated in R (v 4.4) using ggplot2 (117, 118).

### Quantitative RT-PCR and semi-quantitative PCR

DNase-treated RNA was reverse-transcribed with iScript cDNA Synthesis Kit (Bio-Rad). qPCR was performed with PowerUp SYBR Green Master Mix (Thermo Fisher) on a QuantStudio Flex6 instrument. For gel-based analysis, Q5 polymerase (NEB) amplicons were resolved on 2 % agarose and visualized on a Li-COR Odyssey imager.

### Immunoblotting

Cells were lysed in 1 % SDS, boiled, and analyzed by SDS-PAGE (4–20 % gradient) and trans-fer to 0.2 uM PVDF membranes (Immobilon-PSQ Millipore Sigma). Primary antibodies included anti-FLAG (FG4R Invitrogen, 1:1000), anti-IRGM (E6P7W Cell Signaling Technology, 1:1000), anti-HNRNPA2B1 (3H6F7 ProteinTech, 1:1000), and anti-β-actin (ab8226 Abcam, 1:1000). Secondary antibodies included Goat-anti-mouse IgG-800CW (926-32210 Licor, 1:10000) and Goat-anti-rabbit IgG-680RD (926-68071 Licor, 1:10000).

### Immunofluorescence microscopy

For immunofluorescence, cells on coverslips were fixed (4 % PFA), permeabilized (0.2 % Triton X-100), stained with primary and fluorophore-conjugated secondary antibodies, and mounted in ProLong Diamond. Fluorescence microscopy was performed using Zeiss LSM880 Airyscan and LSM710 confocal microscopes through the Vanderbilt Cell Imaging Shared Resource. Primary antibodies included anti-FLAG (FG4R Invitrogen, 1:500), anti-LC3 (L10382 Thermofisher, 1:500), anti-LAMP1 (E5N9Z Cell Signaling Technology, 1:500).

### Image processing and segmentation

Cells were segmented using DIC/BF images and Cellpose (cyto2 model) with standard preprocessing and small-object filtering. Mtb bacilli were identified from the Mtb fluorescence channel by intensity-based thresholding and morphological cleanup.

### Fluorescence Quantification, Colocalization, and Statistics

Recruitment of LC3 and LAMP1 to Mtb was quantified by measuring mean fluorescence intensity within a thin ring surrounding each segmented bacillus, as well as within whole-cell cytoplasmic masks. Bacilli positive for each marker were defined using a uniform intensity threshold applied across conditions. Colocalization between markers and Mtb was assessed using Manders’ (M1/M2) and Pearson correlation coefficients computed from background-subtracted pixel intensities within cytoplasmic or bacillus-associated regions. All statistical analyses were performed in Python using SciPy and custom permutation tests. Group comparisons used appropriate parametric or nonparametric tests with Benjamini–Hochberg correction where indicated. At least 300 cells per condition were analyzed, and error bars represent SD unless noted otherwise.

### Live cell imaging and analysis

Cells were incubated in 100 nm lysotracker-deepred (L12492 Thermofisher) and 1 ug/mL Hoechst stain (H21486 Thermofisher) for 30 minutes then infected with *E. coli* stably expressing mCherry at an MOI = 10 and centrifuged (1000x RPM VWR MegaStar 4.0, 10 minutes) briefly to synchronize infection. Media was replaced with optical complete DMEM supplemented with 1 ug/mL Hoechst stain. Time-lapse CZI files were acquired on a Zeiss LSM880 Airyscan microscope (four channels: LysoTracker, DIC, DAPI, mCherry) using identical settings (0.215-µm pixels; 2-min intervals; 1h duration). Images were Z-projected and segmented in Cellpose to iden-tify nuclei and cytoplasms. For each cell and timepoint, LysoTracker and mCherry mean intensities and Pearson/Manders colocalization coefficients were quantified. Cells were tracked by nearest-centroid linking (5-µm maximum displacement), and per-cell metrics were aggregated across movies to generate genotype-specific lysosomal recruitment and bacterial colocalization dynamics.

### Functional assays

Cells (2.5 × 10⁴/mL) were spin-infected (10 min, 1000 rpm; MOI 10), washed once with PBS, and overlaid with 0.1% PI-containing medium. PI fluorescence was measured on a TECAN Spark plate reader. Triton X-100 (0.1%) wells served as genotype-matched positive controls and media-only wells as background. Percent cell death was calculated as:

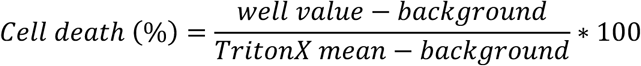

### Statistical analysis and figure generation

All statistical analyses were performed in GraphPad Prism (v 10.4.1). Two-tailed unpaired Student’s t tests or Mann-Whitney tests were applied as appropriate. Data are mean ± SEM from ≥3 biological replicates. Our model was made with BioRender (RI293O6ST8).

### Data availability

RNA-seq data supporting this study are available from the Gene Expression Omnibus (GEO accession GSE313377). Additional data and reagents are available from the corresponding authors upon request.

## ACKNOWLEDGEMENTS

We would like to thank members of Patrick and Watson labs for feedback and experimental help throughout these studies. We’d like to acknowledge the Genomic and RNA Profiling Core (GARP) at Baylor College of Medicine for hnRNPA2B1 KO library preparation and RNA-seq. Thanks to Cindy Nochowicz and the Vanderbilt Tuberculosis Center for support of the BSL3 suite and to the Cell Imaging Shared Resource (CISR) at Vanderbilt School of Medicine for microscopy training and assistance with equipment. Funding was provided by NIH/NIGMS to K.L.P. (R35GM133720), NIH/NIAID to R.O.W. and K.L.P. (R01AI155621), NIH/NIGMS to J.B.H. (5T32GM137793-05), NIH/NIAID to A.K.C. (1F31AI176652-01A1), and NIH/NIAID to C.J.M (5F31AI76795-02).

